# L-DOPA enhances hippocampal direction signals in younger and older adults

**DOI:** 10.1101/2021.08.18.456677

**Authors:** Christoph Koch, Christian Bäuchl, Franka Glöckner, Philipp Riedel, Johannes Petzold, Michael Smolka, Shu-Chen Li, Nicolas W. Schuck

## Abstract

Previous studies indicate a role of dopamine in hippocampus-dependent spatial navigation. Although neural representations of direction are an important aspect of spatial cognition, it is not well understood whether dopamine directly affects these representations, or only impacts other aspects of spatial brain function. Moreover, both dopamine and spatial cognition decline sharply during age, raising the question which effect dopamine has on directional signals in the brain of older adults. To investigate these questions, we used a double-blind cross-over L-DOPA/Placebo intervention design in which 43 younger and 37 older adults navigated in a virtual spatial environment while undergoing functional magnetic resonance imaging (fMRI). We studied the effect of L-DOPA, a DA precursor, on fMRI activation patterns that encode spatial walking directions that have previously been shown to lose specificity with age. This was done in predefined regions of interest, including the early visual cortex, retrosplenial cortex, and hippocampus. Classification of brain activation patterns associated with different walking directions was improved in the hippocampus and the retrosplenial cortex following L-DOPA administration. This suggests that DA enhances the specificity of neural representations of walking direction in these areas. In the hippocampus these results were found in both age groups, while in the RSC they were only observed in younger adults. Taken together, our study provides evidence for a mechanistic link between DA and the specificity of neural responses during spatial navigation.

**Significance Statement:** The sense of direction is an important aspect of spatial navigation, and neural representations of direction can be found throughout a large network of space-related brain regions. But what influences how well these representations track someone’s true direction? Using a double-blind cross-over L-DOPA/Placebo intervention design, we find causal evidence that the neurotransmitter dopamine impacts the fidelity of direction selective neural representations in the human hippocampus and retrosplenial cortex. Interestingly, the effect of L-DOPA was either equally present or even smaller in older adults, despite the well-known age related decline of dopamine. These results provide novel insights into how dopamine shapes the neural representations that underlie spatial navigation.

## 1 Introduction

A role of dopamine (DA) in spatial navigation is well established. Anatomically, spatial cognition depends on a network of brain regions centered around the hippocampus (HC) (Burgess, Maguire, & O’Keefe, 2002; Chersi & Burgess, 2015) that is a target of dopaminergic innervation from the ventral tegmental area and the locus coeruleus (McNamara & Dupret, 2017). Behaviorally, spatial navigation abilities are influenced by DA functioning in younger as well as older animals and humans (Granado et al., 2008; El-Ghundi et al., 1999; Thurm et al., 2016; Kentros, Agnihotri, Streater, Hawkins, & Kandel, 2004).

Much less is known about how DA might change the neural representations that support spatial navigation. Particularly interesting for human neuroscience are direction selective representations (Taube, 2007), which have been found, amongst others, in the HC, the retrosplenial cortex (RSC) and visual cortex (Shine, Valdés-Herrera, Hegarty, & Wolbers, 2016; Flossmann & Rochefort, 2021; Guitchounts, Masís, Wolff, & Cox, 2020; Cacucci,Lever, Wills, Burgess, & O’Keefe, 2004), and can be decoded from human fMRI signals (Koch, Li, Polk, & Schuck, 2020). We hypothesized that DA affects direction encoding in the human brain and tested this idea using a double-blind placebo controlled intervention design. Specifically, we predicted that oral administration of L-DOPA, a dopamine precursor, would influence how accurately walking direction can be decoded from multi-voxel fMRI patterns.

Next to its role in spatial navigation, DA has also received much attention in the context of aging, where reduced DA functions are prevalent and are thought to underlie age-related cognitive declines (Bäckman, Nyberg, Lindenberger, Li, & Farde, 2006; Li, Lindenberger, & Bäckman, 2010; Volkow et al., 1998; Chowdhury et al., 2013). Computational models have shown that declining neuromodulatory effects of DA lead to losses in the signal-to-noise ratio of neural responses (Cohen & Servan-Schreiber, 1992; Servan-Schreiber, Printz, & Cohen, 1990), which in the aging brain can lead to neural representations that are less specific or ”dedifferentiated” (Li, Lindenberger, & Sikström, 2001; Li & Rieckmann, 2014). In line with these models, dedifferentiation has repeatedly been observed in older adults (OA) at the behavioral and neural levels (Park et al., 2004; Carp, Park, Polk, & Park, 2011; Carp, Park, Hebrank, Park, & Polk, 2011; Koch et al., 2020; Kobelt, Sommer, Keresztes,Werkle-Bergner, & Sander, 2021; Li et al., 2004). Neural dedifferentiation, in turn, has been linked to decreased memory performance (Koen, Hauck, & Rugg, 2019; Sommer et al., 2019; St-Laurent, Abdi, Bondad, & Buchsbaum, 2014), establishing an explanatory link between DA, neural representations and cognitive aging.

These roles of DA in spatial navigation and aging might contribute to the pronounced decline in spatial cognition with age (Moffat, 2009; Lester, Moffat, Wiener, Barnes, & Wolbers, 2017; Wolbers, Dudchenko, & Wood, 2014; Schuck, Doeller, Polk, Lindenberger, & Li, 2015), and to the neural dedifferentiation of direction-selective (Koch et al., 2020) and hippocampal signals (Schuck et al., 2015) in the aging brain. Moreover, since the sharp decline of DA with age should lead to lower baseline availability of DA in OA, the effects of DA might be stronger in OA relative to younger adults (YA) – reflecting DA’s inverted-U-shape relation to cognitive performance (Cools & D’Esposito, 2011; Li et al., 2013; Vijayraghavan,Wang, Birnbaum, Williams, & Arnsten, 2007; Li et al., 2010). Indeed, one previous study found age-related effects of the DA receptor agonist bromocriptine on dedifferentiation in the HC (Abdulrahman, Fletcher, Bullmore, & Morcom, 2017). Moreover, HC-dependent episodic memory, spatial navigation, and learning have been found to be affected by genetic polymorphisms related to dopamine D2 receptor availability (COMT Val158Met, C957T CC; Papenberg et al., 2014; Li et al., 2013) or hippocampal function (KIBRA SNP rs17070145; Schuck et al., 2013, 2018) in OA, but not YA. Based on these findings, we therefore also tested whether L-DOPA effects on walking direction decoding would be stronger in OA relative to YA.

Finally, we expected that DA could also influence the shape of population-based tuning functions of direction. Although direction-sensitive cells often have a preferred direction, they also fire in response to non-preferred directions in proportion to their similarity to the preferred direction (Taube, 2007). Hence, encoding of direction information seems to follow a Gaussian tuning function, in particular on a population level (Averbeck, Latham, & Pouget, 2006). Research has also shown that age-related neural dedifferentiation results in increased width of such tuning functions with age (Liang et al., 2010; Leventhal, Wang, Pu, Zhou, & Ma, 2003; Schmolesky, Wang, Pu, & Leventhal, 2000), which we too have reported previously using fMRI (Koch et al., 2020). We therefore also investigated whether L-DOPA has effects on the precision of fMRI-derived tuning functions of direction information and whether such effects may interact with age.

## 2 Materials and Methods

### 2.1 Participants

This study was part of a larger project in which the same participants performed multiple tasks, including a sequential decision making task and a virtual reality spatial memory task inside the scanner and other decision tasks outside of the scanner.

Here, we only report results from the MRI analysis of the VR task described below. Specifically, following our previous publication (Koch et al., 2020), our analyses were specific to neural representations of direction signals during the spatial memory task performed while undergoing fMRI. Other data from the same participants was not within the purview of this study and was therefore not investigated. Data of 102 participants which were recruited for two MRI sessions and randomly assigned to one of the two drug intervention groups (i.e., L-DOPA–Placebo or Placebo–L-DOPA) was available for investigating our research question. Ninety-one of these participants (46 OA, 45 YA) successfully completed both sessions. Four OA were excluded from further analyses because they did not respond in at least a third of the trials in at least one of the two sessions. Furthermore, technical issues during data collection led to incomplete or inaccurate data for three other OA, resulting in an overall exclusion of 7 OA. The main sample therefore consisted of 84 participants, out of which 39 were OA (age 65–75, 7 female) and 45 YA (age 26–35, 16 female). Note that the relatively low number of female OA reflects difficulties in recruitment after the onset of the COVID-19 pandemic.

Decoding analyses of the L-DOPA effects introduced additional requirements for the distribution of walking direction (see Materials and Methods) that were not met for four participants (2 OA, 2 YA). Thus, the final effective sample for these analyses also excluded these participants and comprise of a total of 37 OA (age 65–75, 6 female) and 43 YA (age 26–35, 16 female).

### 2.2 Virtual Reality Task

During each session of fMRI data collection participants had to complete a similar variant of a spatial memory task that was used in previous studies (Schuck et al., 2015; Thurm et al., 2016). Analyses of the present work are mainly concerned with directional signals obtained during free navigation, and hence focus on the corresponding task phases. Specifically, to avoid effects of changed environmental cues on directional signals (e.g.Taube, Muller, & Ranck, 1990) or initial learning, we considered only data from the feedback phase for this study (see below). On average, the included data reflected a period of 17.36 minutes from free navigation per session.

Briefly, participants were placed in a virtual, circular arena in which they could move around freely using a custom-made MRI-compatible joystick. The arena consisted of a circular grass plane surrounded by a wall. Participants could also see distal cues (mountains, clouds) as well as a local cue (traffic cone) to aid orientation (see Fig 1). We asked participants to remember the location of five objects within the 360°arena. First, an initial encoding phase took place in which participants could see and walk to the locations of all objects appearing one after the other. Learning of object location then continued in a feedback phase: participants were placed close to the center of the arena with a random heading direction. After the brief presentation of a grey screen and fixation cross, a picture of the first object was shown. Participants were asked to navigate as closely as possible to the location of this object and indicate their final position with a button press within a maximum of 60 seconds. To provide feedback, the true object location was shown to participants following their response, and they were then asked to navigate to and walk over the shown location. After the feedback, participants were shown another object and the procedure repeated without placing the player in the center of the arena until all five objects were completed. The order in which the five objects were shown was pseudo-randomized. Once all five objects were completed, participants were again placed close to the arena’s center and had to navigate to all five objects in the same manner for a total of six repetitions (i.e., 5 × 6 = 30 feedback trials). In a final transfer phase of the task (data not analyzed in this study, see above), either the arena size or the location of the traffic cone were altered, and participants‘ object location memory was tested again as above. For the second session participants had to learn the location of five different objects, but the trial structure and procedures were identical otherwise. Completing one session took participants between 14 and 49 minutes.

**Figure 1:**
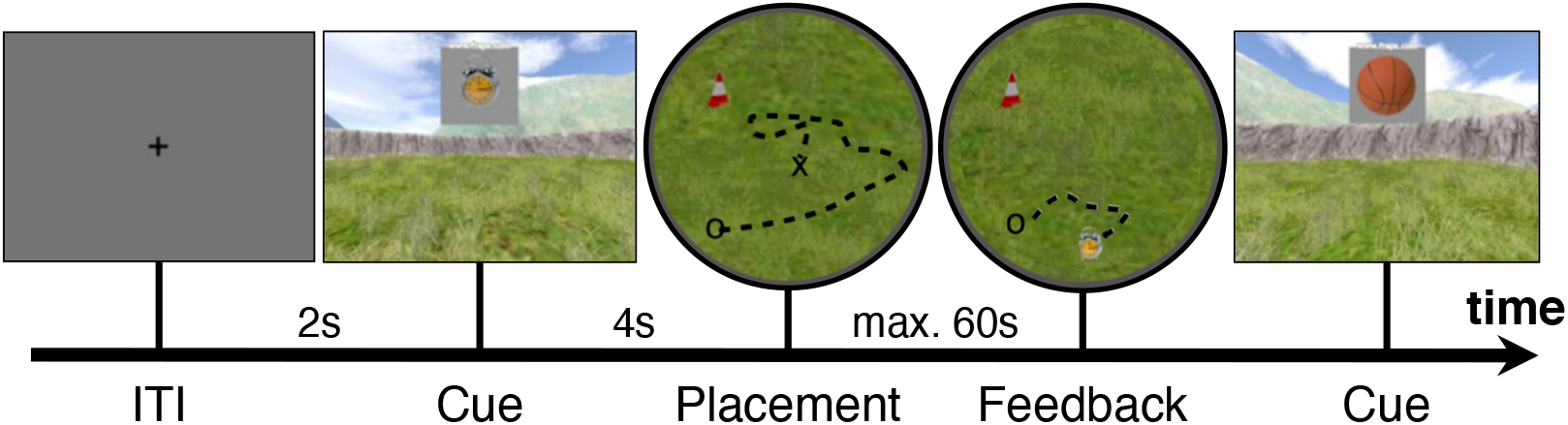
Task procedure during feedback phase. Each trial started with a fixation cross on a grey background for two seconds. Afterwards a cue was presented showing the object to which participants needed to navigate (object locations were learned during encoding phase). The participant then had 60 seconds to navigate from their starting location (cross) to the object location according to their spatial memory. Participants indicated that they had arrived at the remembered location (circle) by pressing a response button, after which the object appeared at its true location. Participants could observe the difference between their response and the correct location and were required to navigate towards and walk over the correct location, before the cue of the next trial was presented.

### 2.3 Drug administration

Following a double-blind drug administration design, participants were given either a total of 225mg of L-DOPA (Madopar, Roche, Levodopa/Benserazid, 4:1 ratio) or a placebo (P-Tabletten white 8mm Lichtenstein, Winthrop Arzneimittel) before each MRI session in the form of two orally administered dosages. A first dosage (150mg L-DOPA/Placebo) was given about 10 minutes before subjects entered the MRI scanner, roughly one hour before the spatial navigation task began. To assure high dopamine availability during the task, a second booster dosage (75mg L-DOPA/Placebo) was administered roughly ten minutes before task onset (cf.Kroemer et al., 2019). Participants were pseudo-randomly assigned to one of two groups with different session order, either the group that received L-DOPA in the first session and placebo in the second session (Drug-Placebo group, 40 subjects) or the group that started with the placebo in the first session (Placebo-Drug group, 44 participants).

### 2.4 Image acquisition

All data was collected on a 3 Tesla Siemens Magnetom Trio (Siemens, Erlangen, Germany) MRI scanner. T1-weighted structural images were collected at the beginning of the first session using a MP-RAGE pulse sequence (0.8 × 0.8 × 0.8 *mm* voxels, TR = 2400 *ms*, TE = 2.19 *ms*, TI = 1000 *ms*, acquisition matrix = 320 × 320 × 240, FOV = 272 *mm*, flip angle = 8°, 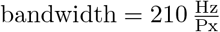). At the beginning of the second session T2-weighted structural scan was collected (0.8 × 0.8 × 0.8 *mm* voxels, TR = 3200 *ms*, TE = 565 *ms*, acquisition matrix = 350 × 350 × 2630, FOV = 272 mm, 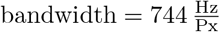).

Functional on-task data was collected using a T2*-weighted echo-planar imaging (EPI) pulse sequence 3 × 3 × 2.5 *mm* voxels, slice thickness = 2.5 *mm*, distance factor = 20%, TR = 2360 *ms*, TE = 25 *ms*, image matrix = 64 × 64, FOV = 192 *mm*, flip angle = 80°, 48 axial slices, GRAPPA parallel imaging, acceleration factor: 2, interleaved acquisition). The sequence lasted until the task was completed and took about 15 – 50 minutes. Additional functional scans not analyzed in this manuscript included data from the transfer phase, data from a decision making task, as well as data from a resting state scan collected at the start of each session.

Quality of all collected functional sequences was assessed using MRI quality control (MRIQC;Esteban et al., 2017). The quality measure of framewise displacement (FD, threshold 3mm), a measure for movement during image acquisition (Power et al., 2014), was extracted and used for statistical control.

### 2.5 ROIs

Each ROI was created from anatomical labels obtained from Mindboggle’s FreeSurfer-based segmentation of each participant’s individual T1-weighted images (Klein et al., 2017). We investigated three predefined ROIs in light of previous findings indicating direction selective coding in these regions (Taube, 2007; Shine et al., 2016; Flossmann & Rochefort, 2021; Guitchounts et al., 2020; Cacucci et al., 2004; Koch et al., 2020). An early visual cortex (EVC) ROI, consisting of the bilateral cortical masks of the cuneus, lateral occipital cortex, and the pericalcarine cortex. A ROI of the retrosplenial cortex (RSC) constructed from the bilateral, cortical masks of the cingulate ishtmus. A mask of the hippocampus (HC) was extracted from the respective bilateral masks of the parcellation. In addition to these core masks, we added a ROI of the left motor cortex, constructed from the cortical mask of the left precentral gyrus, to serve as a control. Although our resolution was suboptimal to investigate small areas, we included a mask of the entorhinal cortex (EC) in order to explore if direction signals could be found there as well.

### 2.6 Image preprocessing

#### Copyright Waiver

Results included in this manuscript come from preprocessing performed using *fMRIPrep* 20.0.6 (Esteban, Markiewicz, et al., 2018; Esteban, Blair, et al., 2018; RRID:SCR_016216), which is based on *Nipype* 1.4.2 (Gorgolewski et al., 2011, 2018; RRID:SCR_002502). The boilerplate text in this section (2.6) was automatically generated by fMRIPrep with the express intention that users should copy and paste this text into their manuscripts *unchanged*. It is released under the [CC0](https://creativecommons.org/publicdomain/zero/1.0/) license.

#### Anatomical data preprocessing

The T1-weighted (T1w) image was corrected for intensity non-uniformity (INU) with `N4BiasFieldCorrection` (Tustison et al., 2010), distributed with ANTs 2.2.0 (Avants, Epstein, Grossman, & Gee, 2008; RRID:SCR_004757), and used as T1w-reference throughout the workflow. The T1w-reference was then skull-stripped with a *Nipype* implementation of the `antsBrainExtraction.sh` workflow (from ANTs), using OASIS30ANTs as target template. Brain tissue segmentation of cerebrospinal fluid (CSF), white-matter (WM) and gray-matter (GM) was performed on the brain-extracted T1w using `fast` (FSL 5.0.9; RRID:SCR_002823;Zhang, Brady, & Smith, 2001). Brain surfaces were reconstructed using `recon-all` (FreeSurfer 6.0.1; RRID:SCR_001847;Dale, Fischl, & Sereno, 1999), and the brain mask estimated previously was refined with a custom variation of the method to reconcile ANTs-derived and FreeSurfer-derived segmentations of the cortical gray-matter of Mindboggle (RRID:SCR_002438;Klein et al., 2017). Volume-based spatial normalization to two standard spaces (MNI152Lin, MNI152NLin2009cAsym) was performed through nonlinear registration with `antsRegistration` (ANTs 2.2.0), using brain-extracted versions of both T1w reference and the T1w template. The following templates were selected for spatial normalization: *Linear ICBM Average Brain (ICBM152) Stereotaxic Registration Model* (Mazziotta, Toga, Evans, Fox, & Lancaster, 1995; TemplateFlow ID: MNI152Lin), *ICBM 152 Nonlinear Asymmetrical template version 2009c* (Fonov, Evans, McKinstry, Almli, & Collins, 2009; RRID:SCR_008796; TemplateFlow ID: MNI152NLin2009cAsym).

#### Functional data preprocessing

For each of the 4 BOLD runs collected per subject (two task related runs reported here and 2 resting state runs not reported here), the following preprocessing was performed. First, a reference volume and its skull-stripped version were generated using a custom methodology of *fMRIPrep*. Susceptibility distortion correction (SDC) was omitted. The BOLD reference was then co-registered to the T1w reference using `bbregister` (FreeSurfer) which implements boundary-based registration (Greve & Fischl, 2009). Co-registration was configured with six degrees of freedom. Head-motion parameters with respect to the BOLD reference (transformation matrices, and six corresponding rotation and translation parameters) are estimated before any spatiotemporal filtering using `mcflirt` (FSL 5.0.9;Jenkinson, Bannister, Brady, & Smith, 2002). BOLD runs were slice-time corrected using `3dTshift` from AFNI 20160207 (Cox & Hyde, 1997; RRID:SCR_005927). The BOLD time-series were resampled onto the following surfaces (FreeSurfer reconstruction nomenclature): *fsnative, fsaverage*. The BOLD time-series (including slice-timing correction when applied) were resampled onto their original, native space by applying the transforms to correct for head-motion. These resampled BOLD time-series will be referred to as *preprocessed BOLD in original space*, or just *preprocessed BOLD*. The BOLD time-series were resampled into standard space, generating a *preprocessed BOLD run in MNI152Lin space*.The first step in this process was that a reference volume and its skull-stripped version were generated using a custom methodology of *fMRIPrep*. Several confounding time-series were calculated based on the *preprocessed BOLD*: framewise displacement (FD), DVARS and three region-wise global signals. FD and DVARS are calculated for each functional run, both using their implementations in *Nipype* (following the definitions by Power et al., 2014). The three global signals are extracted within the CSF, the WM, and the whole-brain masks. Additionally, a set of physiological regressors were extracted to allow for component-based noise correction (*CompCor*; Behzadi, Restom, Liau, & Liu, 2007). Principal components are estimated after high-pass filtering the *preprocessed BOLD* time-series (using a discrete cosine filter with 128s cut-off) for the two *CompCor* variants: temporal (tCompCor) and anatomical (aCompCor). tCompCor components are then calculated from the top 5% variable voxels within a mask covering the subcortical regions. This subcortical mask is obtained by heavily eroding the brain mask, which ensures it does not include cortical GM regions. For aComp-Cor, components are calculated within the intersection of the aforementioned mask and the union of CSF and WM masks calculated in T1w space, after their projection to the native space of each functional run (using the inverse BOLD-to-T1w transformation). Components are also calculated separately within the WM and CSF masks. For each CompCor decomposition, the *k* components with the largest singular values are retained, such that the retained components’ time series are sufficient to explain 50 percent of variance across the nuisance mask (CSF, WM, combined, or temporal). The remaining components are dropped from consideration. The head-motion estimates calculated in the correction step were also placed within the corresponding confounds file. The confound time series derived from head motion estimates and global signals were expanded with the inclusion of temporal derivatives and quadratic terms for each (Satterthwaite et al., 2013). Frames that exceeded a threshold of 0.5 mm FD or 1.5 standardised DVARS were annotated as motion outliers. All resamplings can be performed with *a single interpolation step* by composing all the pertinent transformations (i.e. head-motion transform matrices, susceptibility distortion correction when available, and co-registrations to anatomical and output spaces). Gridded (volumetric) resamplings were performed using `antsApplyTransforms` (ANTs), configured with Lanczos interpolation to minimize the smoothing effects of other kernels (Lanczos, 1964). Non-gridded (surface) resamplings were performed using `mri_vol2surf` (FreeSurfer).

Many internal operations of *fMRIPrep* use *Nilearn* 0.6.2 (RRID:SCR_001362;Abraham et al., 2014), mostly within the functional processing workflow. For more details of the pipeline, see [the section corresponding to workflows in *fMRIPrep*'s documentation](https://fmriprep.readthedocs.io/en/latest/workflows.html).

### 2.7 fMRI analyses

#### Classification of walking direction

All classification of walking direction was performed in Python (Python Software Foundation; Python Language Reference, version 3.7.8; available at http://www.python.org) and relied on `scikit-learn` (Pedregosa et al., 2011) and `nilearn` (Abraham et al., 2014). Statistical analysis was performed using R (version 4.0.3,R Core Team, 2021) and the packages `lme4` (Bates, Mächler, Bolker, & Walker, 2015) and `emmeans` (Lenth, 2021). All statistical figures were created using the `ggplot2` package (Wickham, 2016).

Functional data was prepared for classification by smoothing images with a 3mm FWHM kernel. Next, `nilearn`’s `signal.clean` function was used to detrend, high-pass filter 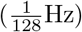, de-noise (using 10 components of aCompCor) and z-standardize the time courses.

Participants’ walking direction was extracted from navigated paths within the virtual environment. The complete 360°-space of direction was binned into six equally spaced bins of 60°. Classifier training examples were then constructed by taking fMRI multi-voxel patterns in response to consistent walking within one binned direction for at least one second. Hence the number of classifier examples for each participant and direction were dependent on the travelled paths and the number of direction changes. If the same example spanned multiple TRs (i.e., was longer than 2.36s) all TRs spanned were averaged to assure a single voxel-pattern per example. Voxel responses were taken two TRs (4.72s) after the event to adjust for hemodynamic lag. A multinomial logistic regression classifier (L2 regularization, C = 1, tolerance = 10^-4^, 1000 maximum iterations; as implemented in `scikit-learn`) was applied to the resulting activation patterns in order to test whether walking direction could be classified. Two cross-validation approaches were used for classification: cross-session and within-session. Cross-session decoding was used to asses overall decoding, irrespective of drug intervention. Within-session decoding was used to separately assess decoding in the L-DOPA and placebo sessions. Results for both approaches are reported separately.

For cross-session cross-validation, in order to reduce auto-correlation of noise, the data of each session was first split into two sets, one consisting of odd and one of even walking direction events. Cross validation approaches as described below were then performed separately for each split. This approach ensured that walking direction events within each of the sets had a higher temporal separation (average: 8.31 seconds, median of 5.70 seconds) as compared to the original data. In consequence, auto-correlation of noise between consecutive examples was reduced, resulting in classifiers that were less biased by autocorrelated event structure (for details, see Koch et al., 2020). Each set was further split into four folds for cross-validation purposes. Specifically, each of the two sessions was split once such that both resulting folds contained the same amount of examples. Separate leave-one-fold-out classification analyses were then performed within each of the two sets (odd/even). The test set, as opposed to the training set, included odd as well as even examples to maximize the number of predictions. Cross-validated decoding results from both sets were combined only afterwards.

Because session was associated with intervention type (placebo or L-DOPA), we also adopted a within-session approach for corss-validation. Specifically, cross-session cross-validation was problematic in two ways: First, it could not be used to asses intervention effects that may differ between sessions. Second, training on data from a DA session and testing on a Placebo session (and vice versa) would risk that DA induced changes in direction specific activation patterns could result in reduced classification. To address these issues, data from one session was separated into three folds, and cross-validated decoding was performed across these folds from the same session. An equal number of events per direction in each fold was ensured as above. The separation into odd and even events was dropped due to reduced data amount when considering only one session. Nevertheless, four participants (2 OA, 2 YA) had to be excluded for missing examples of at least one class in any of the two sessions, leaving a final sample of 80 participants (37 OA, 43 YA).

In both cross-validation approaches, we ensured a balanced number of training examples for each class by upsampling underrepresented classes if necessary. Trained classifiers were then used to predict the walking direction from examples in a testing set given by the remaining fold. A balanced accuracy score was calculated for each test set and results were pooled across all cross-validation runs. The resulting score was compared to a permutation distribution resulting from repeating the same classification 1000 times with randomly permuted class labels in the training set. Additionally, a linear mixed model (LMM) of classification accuracy with fixed effects of age group and ROI, and a random effect of participant was used to asses possible group- or ROI-based differences, as well as their interaction. The model for session-specific decoding results included main- and interaction effects of intervention (L-DOPA vs. Placebo), age group (OA vs. YA), ROI, and session order (L-DOPA – Placebo vs. Placebo – L-DOPA; to allow assessing order effects of the drug intervention). To assess whether drug effects scaled with the administered drug dosage relative to body weight the model also included a relative dosage/kg × intervention interaction. Additionally, in both models main effects of FD and an FD × intervention interaction were included in the model as a nuisance variable to capture possible effects of drug-related head motion. Random effects included a random intercept of participant and a random slope of intervention to assure a within-subject comparison of decoding accuracy in both sessions.

#### Influence of spatial angular difference on fMRI pattern similarity

To test if neural representations of walking direction show the same circular similarity structure as directions in geometrical space, we analysed the structure of classifiers predictions as in Koch et al. (2020). If the similarity of two fMRI patterns of two different directions is associated with their angular distance in space, this should be reflected in the probability distributions over all possible directions. Specifically, we extracted the probability estimates of each of the six classes for each example of the testing set as calculated by the logistic regression classifier. These estimates were aligned with regard to relative angular difference from the target class (−120°, −60°, 0°, 60°, 120°, 180°) and then averaged over all examples, resulting in a single curve for each participant which we refer to as the *confusion function*. Two simple models of the confusion function with one parameter each were compared: A Gaussian curve in the form of

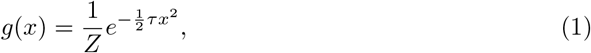

where *x* denotes the angular difference and *τ* the precision (the inverse of the variance, 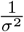). Furthermore, *Z* normalizes the curve. This model captures an inverse relationship between the angular difference of two walking directions and the confusability of their associated neural patterns. An alternative model expressing an absence of such relationship is described by a uniform distribution of classification errors over the remaining five off-center bins. This model could still accommodate high classification for the target class, but would assume that the probabilities of other classes are flat, i.e unrelated to the distance from the target class. Such a model is given by

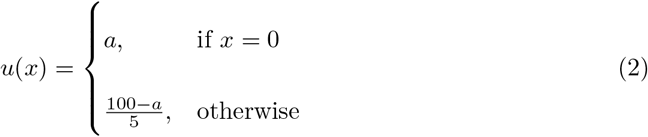

where *a* denotes the classification accuracy. Models were fitted separately within each participant and ROI. Because both models had only one free parameter (*τ* and *a*, respectively), we compared the square root of the mean squared errors (RMSE) between off-center model predictions for both models directly. A better model fit of the Gaussian model indicates directional tuning, i.e. an inverse relationship between the angular difference of directions and the similarity of their neural representations.

In addition to comparing model fits, the Gaussian model allowed us to assess age-differences in directional tuning specificity, which were captured by the precision parameter *τ*. A LMM identical to the one modelling classification accuracy described in the previous section was used to analyze differences in precision.

### 2.8 Behavioral analysis

Task performance during the feedback phase was measured by the distance error: the Euclidean distance between the true location of an object and the location the participant placed the respective object (measured in virtual meters; vm; 1vm = 62.5 Unreal units). Performance for each trial was given by the average distance error across all five presented objects within a trial (missing responses due to exceeding the time limit were excluded). Kolmogorov-Smirnov tests indicated that performance scores of YA were not normally distributed (*D* = .169, *p* = .010, *D* = .064, *p* = .881, for YA and OA, respectively; tested for performance on the last trial). To assure normality, the average distance errors in each trial were log-transformed (*D* = .054, *p* = .941, *D* = .106, *p* = .323 after transform for YA and OA, respectively). To assess the process of learning during the feedback phase of the task, we compared the difference between the first and last trial. Note that in light of non-linear learning curves we did not use a linear model across all trials on purpose. The difference between the two log-transformed measures was modeled using an LMM including the fixed effects of intervention (L-DOPA vs. Placebo), age group, and session order (L-DOPA–Placebo vs. Placebo–L-DOPA) as well as a random intercept of participant. Additionally, we compared performance after learning (last trial) with an identical LMM. Furthermore, group-level performance was compared to chance given by the average distance error assuming random responses for every object. To this end, we uniformly sampled 10^5^ possible locations within the circular arena. The task was then simulated 1000 times while each response of each participant was randomly drawn from the pool of possible locations. This yielded a distribution of 1000 group-means assuming random performance over a given trial and allowed a comparison of trial-specific group-means

Finally, we aimed to quantify the relationship between the specificity of direction signals and task performance to see if more specific direction signals allow better performance on the given task. To this end, we used previous LMMs of classification accuracy but added the regressor of performance in the last trial of the experiment. To assure normally distributed values the log-transformed performance variable was used. Furthermore, performance values were demeaned to eliminate a possible confound between age group and task performance. The FD-related nuisance regressors as well as the interaction between dosage per body weight and intervention were dropped from the model. To see if L-DOPA enhanced signal specificity in proportion to its enhancement of task performance the above model was adapted to predict the difference between sessions in classification accuracy (L-DOPA - Placebo). The increase in task performance was given by the session difference (L-DOPA - Placebo) of the log-transformed performance in the last trial of the task.

## 3 Results

### 3.1 Behavioral results

Log-transformed average distance errors on each trial for both age groups and interventions (L-DOPA vs. placebo) are displayed in Fig. 2. We first investigated log transformed distance errors on the last trial after learning, using a linear mixed model with fixed effects of interest for intervention and age group and a random effect of participant.

**Figure 2:**
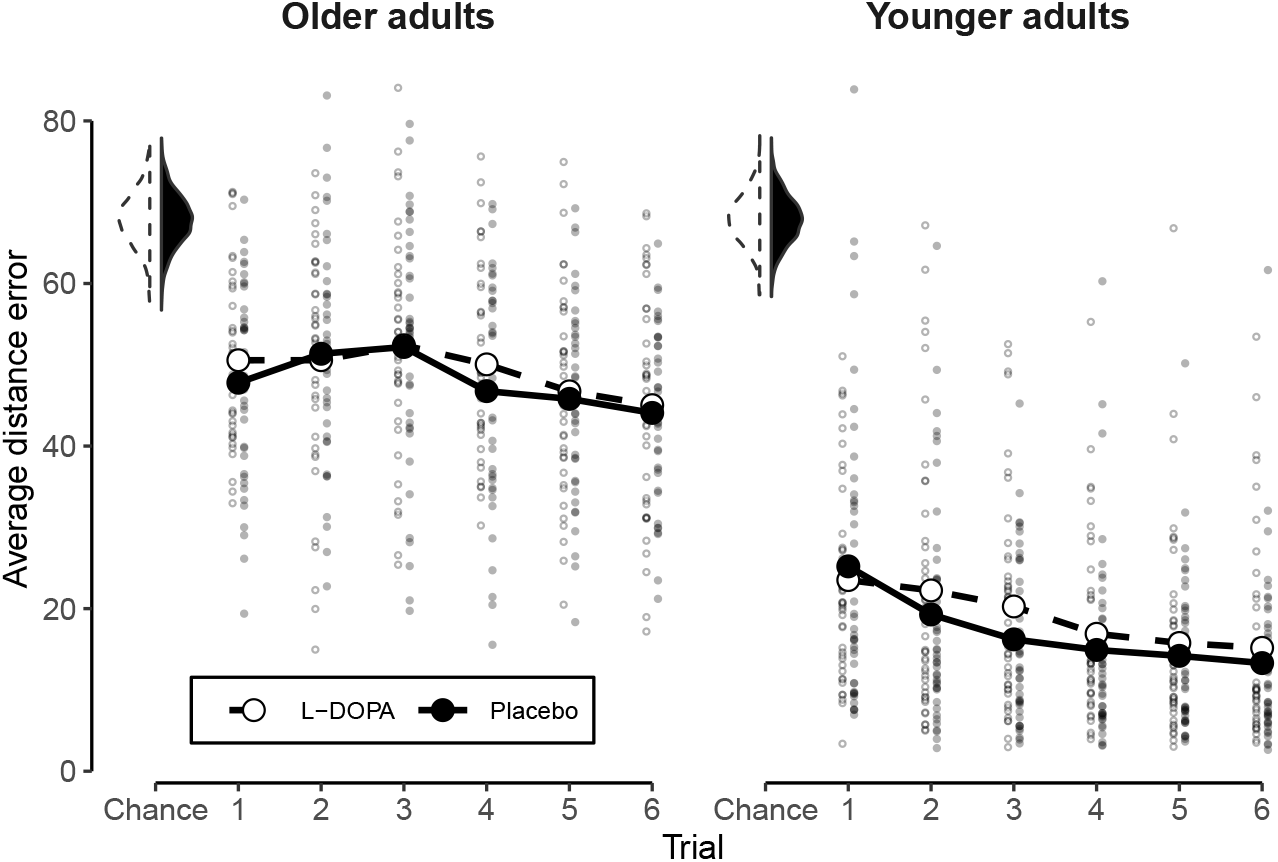
Behavioral results. Average error in object placement for all six trials for OA and YA. Error was measured as the Euclidean distance in vm between the true location of an object and the participants’ placement. Reduction in error shows better task performance. All values of the placebo session depicted in black, all values of the L-DOPA session depicted in white. Small dots indicate individual values of participants. Average over participants in each trial shown by the large dots. Shown on the upper left are session-specific distributions of 10^3^ average performance values in a trial assuming random placement of objects. Note that, in turn, only the trial averages (large dots) can be compared to this chance-distribution.

The LMM showed a significant main effect of age group (*χ*^2^(1) = 167.010, *p* > .001, the *χ*^2^ reflect likelihood ratio tests, see Methods). Post-hoc tests showed that OA had higher distance errors compared to YA at the end of learning (*t*(80) = 12.811, *p* < .001). The model did not display any significant main effect of intervention (*χ*^2^(1) =1.479, *p* = .224) or intervention by age interaction.

Next, we investigated performance increases, i.e. log distance errors on the first minus the last trial. Again, a LMM revealed a significant main effect of age group (*χ*^2^(1) = 61.054, *p* > .001), but no main effect of intervention or intervention × age interaction.

Investigating the nuisance variable of session order revealed no main effects in either end-of-learning performance (*χ*^2^(1) = 0.1784, *p* = .673) or in performance changes (*χ*^2^(1) = 0.948, *p* = .330). No session order × intervention effect was found for performance changes. Unexpectedly, we found a significant interaction of intervention × session order in end-of-learning performance (*χ*^2^(1) = 13.744, *p* < .001), reflecting a trend for a positive effect of DA if L-DOPA was given in the second session (*t*(80) = −1.693, *p* = .094), while this was reversed if L-DOPA was given in the first session (*t*(80) = 3.368, *p* = .001).

### 3.2 Decodability of walking direction

We first assessed decoding using a cross-validation approach across intervention sessions (see Methods). In line with previous work (Koch et al., 2020), walking direction could be decoded in the EVC (*p* < .001) and RSC (*p* = .040), but not in the left motor cortex (*p* = .255), entorhinal cortex (*p* > .999) and HC (*p* > .999), compared to a permutation test (Bonferroni corrected for five comparisons, one sided). A LMM of classification accuracy with fixed effects of interest for age group and ROI, and a random effect of participant revealed a significant main effect of age group (*χ*^2^(1) = 16.209, *p* < .001), a main effect of ROI (*χ*^2^(4) = 194.810, *p* < .001), and a ROI × age group interaction (*χ*^2^(4) = 37.851, *p* > . 001). Post-hoc mean comparisons showed significantly higher classification accuracy in YA in the EVC (*t*(320) = −7.280, *p* < .001), but no such age-related effects in the RSC or HC (*t*(320) ≥ −1.489, *p* ≥ .138), see Fig. 3A. Although classification accuracy was higher in YA in the EVC, a permutation test revealed significant above-chance decoding also in OA (*p* < .001). No main effect of the nuisance variable FD was found (*χ*^2^(1) = .482, *p* > .487).

**Figure 3:**
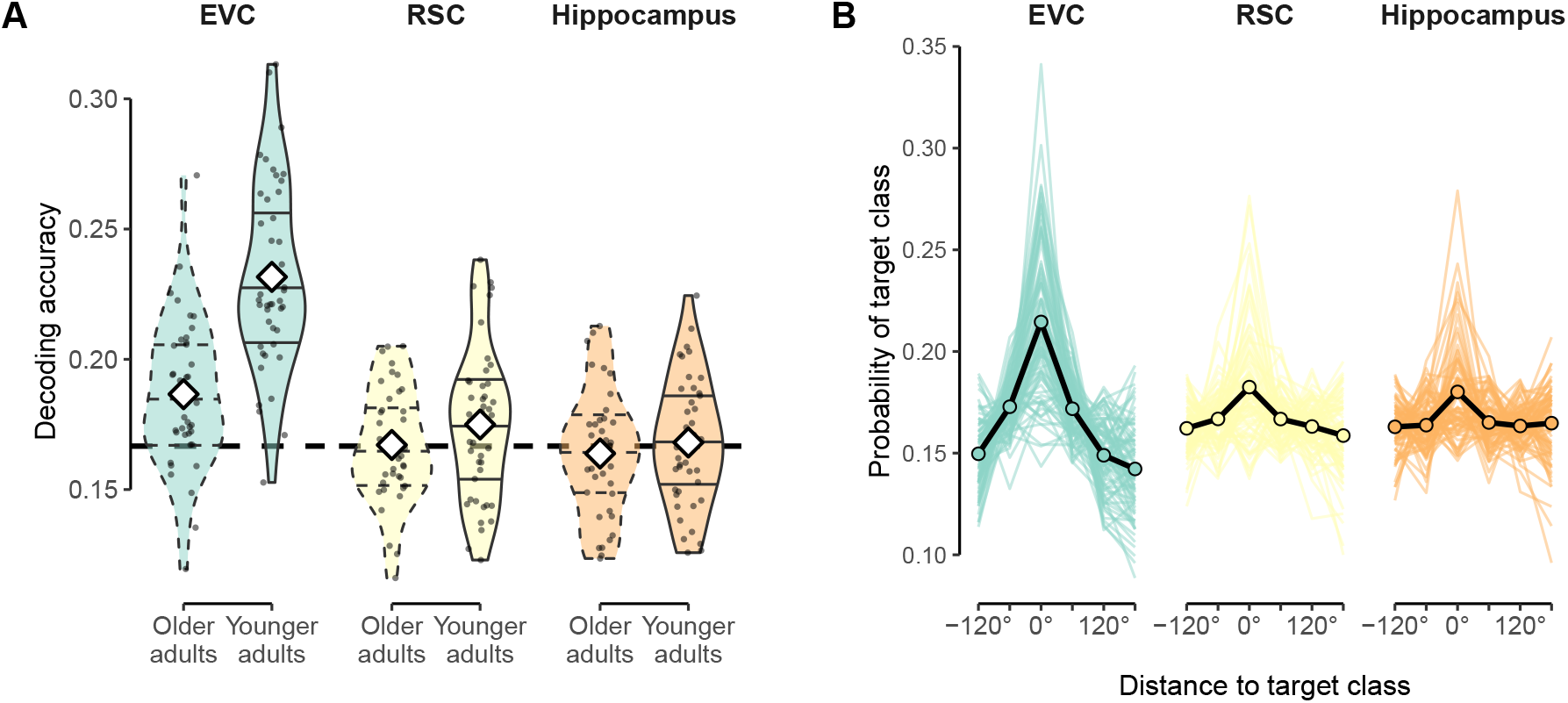
Cross-intervention walking direction decoding. **A**: Cross-validated decoding accuracy of walking direction within both age groups for the cross-session approach, in which the classifier was partially trained on the placebo session and tested on the DA session, or vice versa. Results are shown separately for EVC (green), RSC (yellow), and hippocampus (orange) and each age group (dashed/solid lines). Violin plots indicate distributions, dots represent individual participants and white diamonds mean accuracy. Horizontal, dashed line indicates chance level. **B**: Confusion function for each classifier. Depicted are class probabilities of the logistic classifier as a function of angular distance from the target class. Colored lines indicate individual participants, black lines the group average. Colors as in A.

Investigating the predicted probabilities by the logistic regression directly, rather than the percent of correctly predicted events, revealed a peak at the true direction and decreasing probabilities for the off-target directions in RSC and EVC, as expected (see Fig. 3B.). Notably, in this more sensitive analysis also the HC exhibited an above-chance probability of the target direction (*t*(83) = 5.346, *p* < .001, corr.).

### 3.3 Influence of L-DOPA intervention on decodability

Decoding analyses reported above combined data across sessions/interventions and thus cannot be used to examine the effects of intervention type. We therefore used a within-session decoding analysis to investigate the influence of L-DOPA administration on the decodability of walking direction (see Methods). A LMM of classification accuracy indeed showed a significant main effect of L-DOPA intervention (*χ*^2^(1) = 6.796, *p* = .009), which indicated higher decoding in L-DOPA compared to placebo sessions. As before, we also found main effects of ROI (*χ*^2^(4) = 271.674, *p* < .001), but no L-DOPA × ROI interaction (*χ*^2^(4) = 3.847, *p* = .427). Despite the lack of an interaction, post-hoc tests revealed that significantly higher decoding accuracy in the drug compared to the placebo condition was most apparent in the HC (*t*(603) = 2.153, *p* = .032) and trending in the RSC (*t*(603) = 1.916, *p* = .055), while no comparable effects were found in the EVC (*t*(603) = 1.447, *p* = .148). Results are displayed in Fig. 4A.

**Figure 4:**
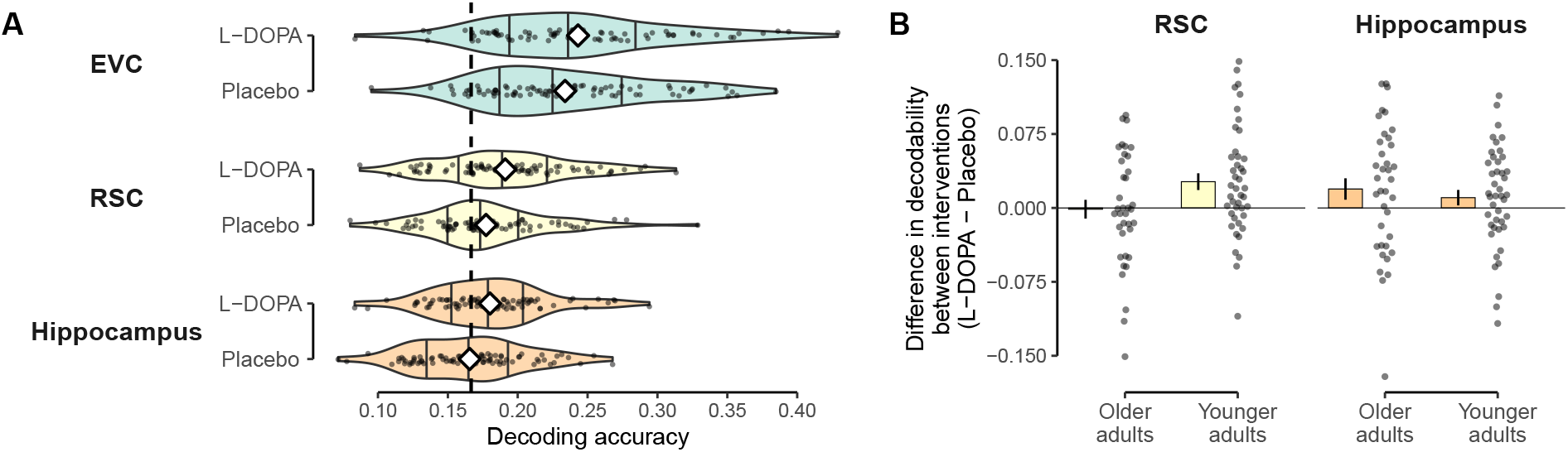
Effect of L-DOPA on decoding of neural walking direction signals. **A**: Intervention-specific decodability of walking direction within each ROI. Black dots show values of participants and violin plots depict intervention-specific distribution. Means are represented by white diamonds. Chance-level is shown by dashed line and based on the total number of classes (6 classes, 16.6% chance). **B**: Influence of drug intervention on decodability (L-DOPA – Placebo) shown for the RSC and hippocampus and split by age groups. Values higher than zero indicate higher decoding accuracy in the L-DOPA condition. Bars reflect group means and error bars reflect SEM. Black dots show individual values of each participant.

In addition, we also found a main effect of age group (*χ*^2^(1) = 6.273, *p* = .012) and a age group × ROI interaction (*χ*^2^(4) = 60.970, *p* < .001). We will elucidate these age effects further below, using separate LMMs per ROI. None of the included nuisance regressors of FD (*χ*^2^(1) = 3.064, *p* = .080) or an interaction between FD and intervention (*χ*^2^(1) = .048, *p* = .826) showed a significant effect on decodability of walking direction. The same was true for any effects of session order (*χ*^2^(1) = .083, *p* = .774). There was no significant interaction between the intervention and the administered dosage per body weight (*χ*^2^(2) = .286, *p* = .867).

To further specify the region-specific effects of DA, LMMs were run separately for each ROI. These ROI-specific LMMs reproduced the main effects of intervention within the HC (*χ*^2^(1) = 5.263, *p* = .022) and the RSC (*χ*^2^(1) = 4.868, *p* = .027). In addition, we found an intervention × age group interaction within the RSC (*χ*^2^(1) = 3.877, *p* = .049), but no such interaction in HC (*χ*^2^(1) = 1.518, *p* = .218). Post-hoc comparisons showed that the effect in RSC was driven by higher decodability of walking direction in the DA compared to placebo session in young adults (*t*(75.6) = 2.879, *p* = .005), but not in OA (*t*(75.4) = −.161, *p* = .872). Within the EVC, only a main effect of age group (*χ*^2^(1) = 16.350, *p* < .001), but no effect of DA intervention (*χ*^2^(1) = 2.038, *p* = .153) was found.

Fig. 4B shows the increase in decodability of walking direction in the L-DOPA condition for the HC and RSC, respectively for each age group. Note that the random slope of intervention had to be dropped from these models to avoid having the same number of random effects as there are data points.

Investigating nuisance variables, we found no impact of dosage per body weight on the intervention effect in any ROI (*χ*^2^(2) < 3.578, *p* ≥ .167, for the interaction). Investigating the movement related variable FD, we found no significant main effects of FD (*χ*^2^(1) ≤ 1.448, *p* ≥ .229) or an interaction between FD and intervention (*χ*^2^(1) ≤ .644, *p* ≥ .422) in HC or RSC. A significant main effect of FD was found in the EVC, however (*χ*^2^(1) = 4.935, *p* = .026). This reflected worse classification accuracy with higher movement during image acquisition (linear regression relating classification accuracy to FD: *b* = −.118, *t*(158) = −6.302, *p* < .001).

A final control analysis within the left motor cortex did neither identify a main effect of intervention (*χ*^2^(1) = .027, *p* = .869) nor any other main effects. Post-hoc tests confirmed that direction decodability under L-DOPA was not significantly different from decodability under placebo, regardless of session order (*t*(74.9) = −1.519, *p* = .133, and *t*(74.1) = 1.202, *p* = .233, L-DOPA–Placebo and Placebo–L-DOPA, respectively).

### 3.4 Influence of L-DOPA intervention on tuning specificity

We next investigated tuning width. Omnibus analyses across ROIs revealed no L-DOPA effect, a main effect of ROI (*χ*^2^(2) = 281.509, *p* < .001), and results otherwise consistent with those reported below. We therefore immediately report results of ROI-specific LMMs.

A model of EVC tuning width found no main effect of intervention or intervention × age effect was found in EVC. We did find a significant main effect of age group (*χ*^2^(1) = 20.631, *p* < .001), reflecting lower precision of the fitted Gaussian curves in OA compared to YA (*t*(79.7) = −4.533, *p* < .001). The same analyses in RSC an HC showed no significant main effects of intervention, age, or intervention × age interactions.

No nuisance effect of FD or FD × intervention interaction were found in any ROI-specific model (*χ*^2^(1) ≤ .857, *p* ≥ .355 and *χ*^2^(1) ≤ .578, *p* ≥ .447, respectively) just as there were no main effects of session order (*χ*^2^(1) ≤ .257, *p* ≥ .612) Additionally, intervention was not involved in any interaction with dosage per body weight (*χ*^2^(2) ≤ 4.412, *p* ≥ .110). Unexpectedly, however, we found a significant intervention × session order interaction in the EVC (*χ*^2^(1) = 10.713, *p* < .001; see Fig. 5A), suggesting that tuning precision was higher when L-DOPA was administered in the second session (*t*(74.0) = 2.911, *p* < .005) compared to when it was administered in the first session (*t*(75.2) = −1.607, *p* = .112). No intervention × session order interaction was found in any other ROI. An exploratory follow up of three-way interactions found a intervention × age group × session order effect in the RSC (*χ*^2^(1) = 6.626, *p* = .010), which pointed towards L-DOPA effects only when given in the second session, and only in YA (*t*(74.6) = 2.818, *p* = .006).

**Figure 5:**
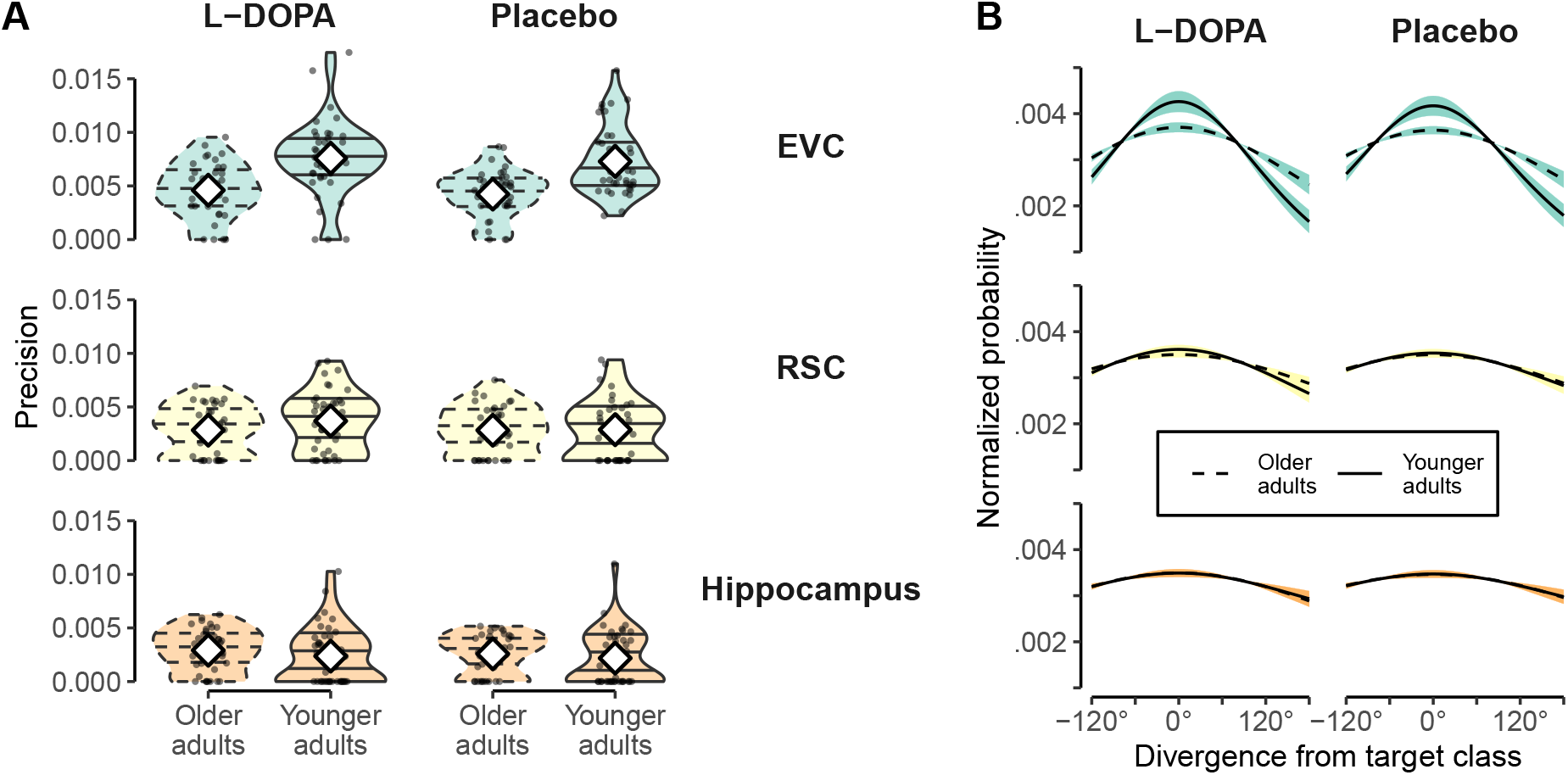
Effect of L-DOPA on tuning specificity. **A**: Precision of Gaussian curves fitted to individual confusion functions in both age groups. Shown separately for the L-DOPA and Placebo intervention in the EVC, RSC, and Hippocampus. Black dots show values of individual participants. Intervention-specific distributions are shown by violin plots. White diamonds depict means. Plots of OA shown in dashed lines for easier distinction. **B**: Mean Gaussian tuning curves shown separately for age groups and intervention (L-DOPA vs. Placebo). ROI separation identical to that of panel A. OA are depicted with dashed lines. Shaded area represents SEM and is colored according to ROI. For each participant a Gaussian curve was fitted to the individual confusion function (given by the classifier). The shown mean Gaussian curves were obtained by averaging participants’ individual Gaussian curves.

The means of the fitted Gaussian curves in the L-DOPA condition are shown in Fig. 5B. Please note that the interpretability of these results is limited since a model comparison between a Gaussian and uniform model of the confusion function remained inconclusive towards either model in both, the drug and placebo condition (*t*(79) ≤ 1.749 or *t*(79) ≥ −1.921, all *p* ≥ .350, corr.).

### 3.5 Relations between task performance, L-DOPA and direction decoding

Finally, we asked whether task performance (spatial distance error) was related to neural direction encoding as well as to the effects of L-DOPA on these neural signals. We therefore investigated the link between session-specific decoding accuracy and task performance on the last trial, in addition to age group, intervention and session order. Because performance on the last trial was highly confounded with age group (see 2) performance values were demeaned within each age group to investigate effects unrelated to age-specific performance differences.

No effects related to task performance were found in the RSC or the HC (*p*s ≥ .053). A model within the EVC revealed a significant main effect of distance error on the last trial on direction decoding (*χ*^2^(1) = 7.594, *p* = .006; see Fig. 6A), pointing towards better decoding accuracy with better task performance (*b* = .040). Besides the main effect, task performance in the EVC also interacted with age group (*χ*^2^(1) = 3.921, *p* = .048), reflecting that the above mentioned relationship was present in YA (*F*(1, 111.03) = 11.912, *p* < .001, *b* = .033) and absent in OA (*F*(1, 121.83) = .066, *p* = .798, *b* = .006). While there was no main effect of session order (*χ*^2^(1) = .009, *p* = .922), the model furthermore indicated a separate interaction between task performance and session order (*χ*^2^(1) = 4.332, *p* = .037). A post-hoc test revealed a trend towards differing slopes depending if L-DOPA was given in the first or second session (*t*(132) = 1.904, *p* = .059) but separate tests within each session order did not display any significant relationships between performance and classification accuracy (*F*(1, 143.83) = .607, *p* = .437, *F*(1, 118.80) = 3.164, *p* = .078, for L-DOPA – Placebo and Placebo – L-DOPA, respectively). As expected the model of EVC decoding accuracy also displayed a main effect of age group (*χ*^2^(1) = 40.244, *p* < .001; see results for influence of DA on decoding accuracy).

**Figure 6:**
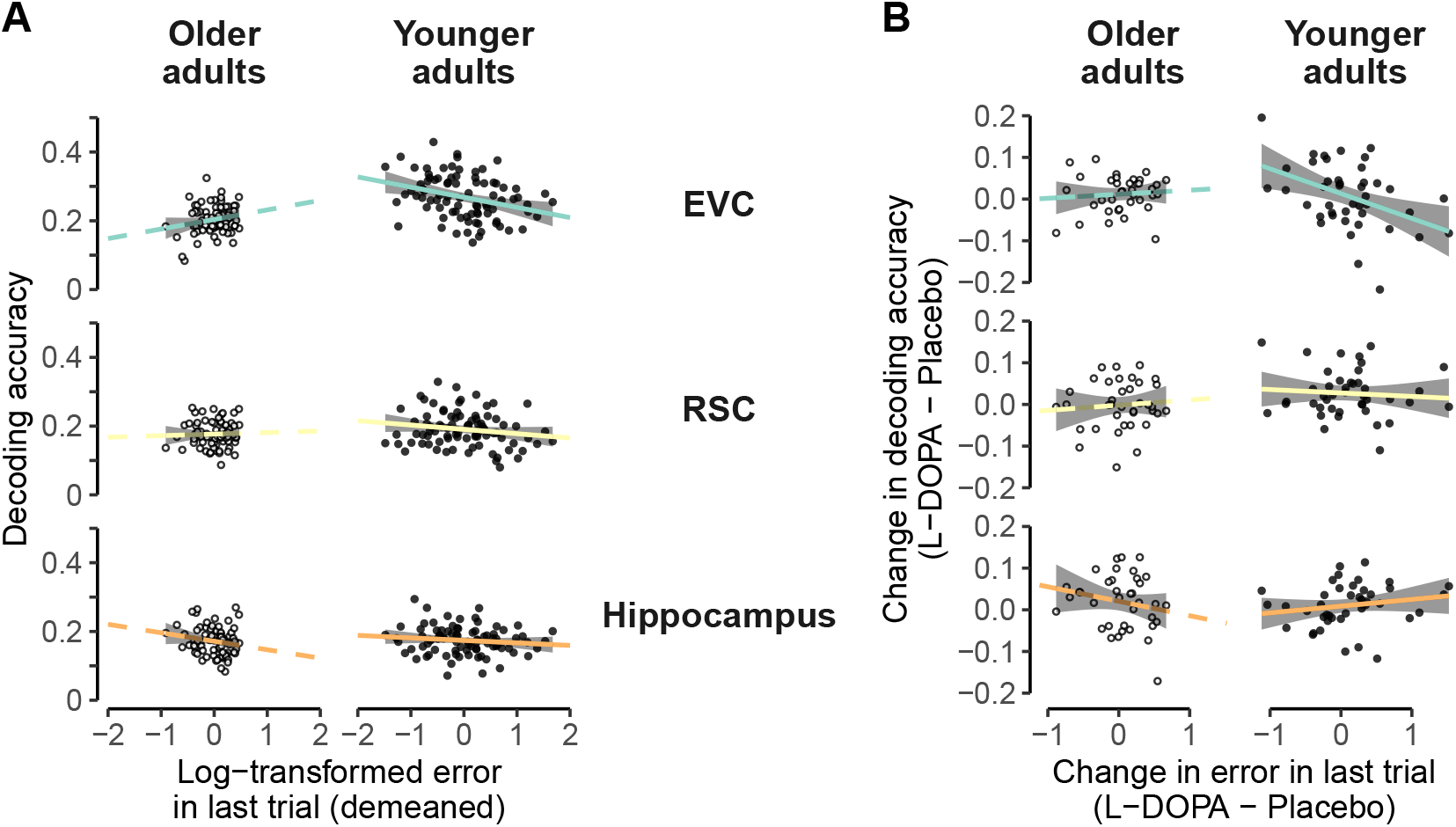
Relationship between decoding accuracy and behavioral performance. **A**: Relationship between decoding accuracy and log-transformed and demeaned distance errors. Shown for the EVC, RSC, and hippocampus separately for both age groups. Dots represent individual participants where OA are shown in white. Lines represent linear models of represented subset and are colored according to the ROI and shown in dashed for OA. **B**: Drug-related change-change relationship between decoding accuracy and behavioral performance. Axes show influence of L-DOPA administration by showing the difference in values between the L-DOPA session and placebo session. Depiction accordingly to A. Please note that in both, A and B, the slope lines were extended beyond the data points purely to aid visibility.

We next investigated change-change relations, asking whether L-DOPA-related changes in decoding were related to L-DOPA-related changes in task performance (see Fig. 6B). Linear regressions revealed that in YA L-DOPA-related changes in direction decoding in EVC were indeed positively related to changes in task performance (*F*(1, 72) = 6.730, *p* = .011, *b* = −.053, negative slopes since performance increase means less errors). In OA this was not the case (*F*(1, 72) = .049, *p* = 826, *b* = .006). Linear models within the RSC and HC did not show any significant effects in change-change relations.

## 4 Discussion

In this work we tested the impact of L-DOPA on neural representations of walking direction in younger and older adults, using a double-blind, cross-over intervention design. In addition to a classic decoding approach, we assessed direction specificity of neural signals, a proxy for tuning functions, using the relative structure of classifier probability estimates. Our results revealed that decodability of walking direction signals in the hippocampus and the retrosplenial cortex was enhanced following the administration of L-DOPA. L-DOPA had comparable effects on HC walking direction signals in both age groups, but in the RSC these DA effects were present only in YA. No L-DOPA effects were found in visual cortex (EVC). Yet, behavioral investigations showed that in younger adults, EVC direction decoding was related to task performance (spatial distance error), and that L-DOPA related changes in EVC decoding were related to changes in task performance. An investigation of tuning specificity revealed no main effects of L-DOPA or L-DOPA × age group interactions.

Furthermore, decoding across interventions, we found evidence for stable direction signals in EVC and RSC, and so some extent also in HPC. Investigating age group differences, we found higher classification accuracy and precision of tuning functions in the EVC of YA compared to OA, a sign of neural dedifferentiation. No age effects on decoding in the HPC or RSC were found. These results confirm our previous finding that neural representations of walking direction can be found in EVC and RSC, and that strong age-related differentiation is present particularly in EVC (Koch et al., 2020). We also showed that better EVC classification accuracy was related to better performance on task, suggesting an important functional role of this area in our task.

Importantly, our results also offer a number of novel insights. First, we show a causal influence of L-DOPA on how walking directions are encoded in the brain, in particular in the HC and the RSC. Both areas have been linked to directional and other spatial information (Spiers & Barry, 2015; Shine et al., 2016; Burles, Slone, & Iaria, 2017), and have even been shown to be part of the same dorsal pathway involved in visuospatial processing (Kravitz,Saleem, Baker, & Mishkin, 2011). Additionally, both areas display dopaminergic innervation (Berger, Verney, Alvarez, Vigny, & Helle, 1985; McNamara & Dupret, 2017), and previous reports have linked DA and spatial cognition more generally (Granado et al., 2008; El-Ghundi et al., 1999; Thurm et al., 2016). Second, the positive effects of DA on decoding are in line with computational models and empirical findings which suggest that DA affects neuronal gain (Li & Rieckmann, 2014; Cohen & Servan-Schreiber, 1992; Thurley, Senn, & Lüscher, 2008). Accordingly, DA’s influence on neural gain could lead to a stronger separation between signal and noise, which made different stimuli more specific and easier to distinguish for the classifier. It should be noted, however, that we did not find any direct effects of L-DOPA on neural direction tuning specificity, which measures how similar neural patterns are to similar directions. Given the effects of DA on neural gain, we had hypothesized that this measure could be more sensitive to the effects of our intervention, but this was not the case. One possible explanation is that our design lacked the power to fully capture the neural tuning functions within just one session. Tentative analyses of EVC and RSC tuning specificity did show DA-related enhancement only in participants who received L-DOPA in the second session. We will discuss these session-specific effects further below.

Third, our study was set up to ask whether the L-DOPA intervention might reduce age-related neural dedifferentiation. Virtual walking direction offered a promising window to answer these questions since it has previously been shown to be subject to age-related neural dedifferentiation (Koch et al., 2020) and the broader domain of spatial cognition has been shown to be highly age-sensitive (Wolbers et al., 2014; Lester et al., 2017). Age is also known to cause substantial loss of DA functioning (e.g. Bäckman et al., 2006), and we speculated that a lower baseline DA availability might magnify the effects of L-DOPA. Surprisingly, we did not find that the effects of L-DOPA were particularly pronounced in OA. Rather, the HC showed age-equivalent effects, and decoding in RSC was in fact enhanced only in YA. Other than individual differences in baseline DA level, task demand may also affect the inverted-U function of DA modulation (Cools & D’Esposito, 2011). The spatial navigation task used in our study is quite demanding, such that YA though have higher baseline DA level could still benefit from the L-DOPA intervention, whereas in OA the task demand may still outweigh the benefit of L-DOPA intervention. While unexpected, these results could offer interesting insights into the complexity of how external DA medication might interact with neural differentiation and compensatory plasticity mechanisms that counteract age-related losses. One notable aspect in this regard is that we found no evidence of age-related dedifferentiation in HC or RSC, which speculatively could be a sign of compensatory mechanisms. It seems possible that DA interventions might only recover neural specificity in brain areas that are affected by age-related dedifferentiation. Contrary to this idea, we found no age-related L-DOPA effects in visual cortex, where dedifferentiation was observed – but this might be due to the relatively low D2 receptor density in this area (Lidow, Goldman-Rakic, Rakic, & Innis, 1989). Another possibility is that we did not observe age-specific effects of L-DOPA on neural direction encoding in RSC and HC for the same reasons we did not find age-related dedifferentiation in these regions. According to this idea, compensatory factors that have mitigated dedifferentiation also affected the effectiveness of external dopamine administration, for instance because of changed connectivity. Both ideas remain speculative and further studies are needed to fully understand how the effects of L-DOPA interventions on neural direction encoding interact with age and dedifferentiation.

Beyond these main implications, a number of interesting observation arose that warrant further investigation. Although we did not find any main effects of session order, we found some indications that session order could influence the effect of L-DOPA on neural signals that underlie spatial navigation. Age-differences in learning were stronger when L-DOPA was administered in the second compared to the first session. In addition, we found tuning specificity in EVC and RSC to be enhanced by L-DOPA only in participants who received the drug in the second session. Stronger effects when DA is administered in a second session have previously been reported in the context of spatial navigation (Thurm et al., 2016). The reason why session order effects could exist in this context are numerous.Garrett et al. (2015), for instance, highlight two possible explanations in the context of DA effects on neural signal variability. One is that previous training may increase the amount of baseline DA-release, based on findings in rodents (Owesson-White, Cheer, Beyene, Carelli, & Wightman, 2008). A DA intervention could therefore lead to differing DA-availability depending on whether the participants had already been trained with the same or a similar task. A second possible explanation raised by Garrett et al. (2015) is that the environment is either learned in a state of higher or normal DA-availability. The state of the second sessions will consequently always be mismatched to the first session, leading to effects of drug administration given the respective session. Related to the first idea, we speculate that in our case general learning about the environment in a first placebo session could have established beneficial baseline for the effects of L-DOPA in the second session. Unfortunately, the present design is unfit to address such explanations and further evidence is warranted.

One open question is why the effect of L-DOPA on decoding in HC and RSC was not reflected in task performance, where no L-DOPA effect was found. In addition to generally small effects on neural representations, another explanation might be that task performance did does only depend on direction signals, but also replies on distance estimation and using distal and local cues, processes which themselves are affected by age (Schuck et al., 2015). The task might therefore have been to complex to provide a suitable behavioral measure. Interestingly, however, we we did find some relationships between behavior and the specificity of directional information in visual cortex, indicating that neural markers might have different relations to performance in our task. This is shown by some of our results also offer insights about age-related changes in the context of spatial navigation more generally. The results in the EVC showed that OA exhibit lower precision of directional tuning functions. This is a replication of findings reported in an earlier study using a similar analysis approach (Koch et al., 2020). During natural navigation and the perception of direction vision plays a major role as it allows stable directional signals (Goodridge, 1998) and corrects and prevents the accumulation of errors during path integration (Jeffery, 2007). A less precise visual signal in OA could therefore influence spatial signals downstream and contribute towards the pronounced difficulties OA have in spatial tasks. Interestingly, we also found a relationship between EVC direction decoding in YA and performance on task, suggesting better spatial memory performance if walking direction could be decoded with higher accuracy. While this concurs with previous reports of a link between (non-spatial) memory and signal specificity (Koen et al., 2019; Sommer et al., 2019; St-Laurent et al., 2014), previous studies have mostly reported such links in older adults. Future work is required to further understand how age-related loss in specificity of visual signals might be involved in spatial cognition. That said, a simple propagation of less specific visual signals to the resplenial complex network seems unlikely, since there was no evidence for age-related dedifferentiation in the RSC or HC.

In summary, we provide first causal insights into the role of dopamine in the encoding of spatial direction signals in the human hippocampus. In addition, our findings show that dopamine also enhances direction encoding in retrosplenial cortex, albeit exclusively in younger adults. In combination with the replication of our own previous results (Koch et al., 2020), these findings offer insights into the neural processes underlying spatial navigation in the human brain, and how they are affected by age more generally.

## Code availability

TBD

## Acknowledgement

The study was supported by a DFG grant to SCL, FG, and MS (SFB 940-2/940-3). NWS was supported by an Independent Max Planck Research Group grant (M.TN.A.BILD0004) awarded by the Max Planck Society and a Starting Grant from the European Union (ERC-2019-StG REPLAY-852669). We want to thank Lorenz Gönner for his helpful comments on the manuscript.

## Conflict of interest

The authors declare no conflicts of interest.

## Notes

### Competing Interest Statement

The authors have declared no competing interest.

## References

Abdulrahman, H., Fletcher, P. C., Bullmore, E., & Morcom, A. M. (2017). Dopamine and memory dedifferentiation in aging. NeuroImage, 153, 211–220. doi: 10.1016/j.neuroimage.2015.03.031

Abraham, A., Pedregosa, F., Eickenberg, M., Gervais, P., Mueller, A., Kossaifi, J., … Varoquaux, G. (2014). Machine learning for neuroimaging with scikit-learn. Frontiers in Neuroinformatics, 8. doi: 10.3389/fninf.2014.00014

Avants, B., Epstein, C., Grossman, M., & Gee, J. (2008). Symmetric diffeomorphic image registration with cross-correlation: Evaluating automated labeling of elderly and neurodegenerative brain. Medical Image Analysis, 12(1), 26–41. doi: 10.1016/j.media.2007.06.004

Averbeck, B. B., Latham, P. E., & Pouget, A. (2006). Neural correlations, population coding and computation (Vol. 7) (No. 5). doi: 10.1038/nrn1888

Bäckman, L., Nyberg, L., Lindenberger, U., Li, S. C., & Farde, L. (2006). The correlative triad among aging, dopamine, and cognition: Current status and future prospects. Neuroscience and Biobehavioral Reviews, 30 (6), 791–807. doi: 10.1016/j.neubiorev.2006.06.005

Bates, D., Mächler, M., Bolker, B., & Walker, S. (2015). Fitting Linear Mixed-Effects Models Using lme4. Journal of Statistical Software, 67(1), 1–48. doi: 10.18637/JSS.V067.I01

Behzadi, Y., Restom, K., Liau, J., & Liu, T. T. (2007). A component based noise correction method (CompCor) for BOLD and perfusion based fmri. NeuroImage, 37(1), 90–101. doi: 10.1016/j.neuroimage.2007.04.042

Berger, B., Verney, C., Alvarez, C., Vigny, A., & Helle, K. B. (1985). New dopaminergic terminal fields in the motor, visual (area 18b) and retrosplenial cortex in the young and adult rat. Immunocytochemical and catecholamine histochemical analyses. Neuroscience, 15(4), 983–998. doi: 10.1016/0306-4522(85)90248-9

Burgess, N., Maguire, E. A., & O’Keefe, J. (2002). The human hippocampus and spatial and episodic memory (Vol. 35) (No. 4). Cell Press. doi: 10.1016/S0896-6273(02)00830-9

Burles, F., Slone, E., & Iaria, G. (2017). Dorso-medial and ventro-lateral functional specialization of the human retrosplenial complex in spatial updating and orienting. Brain Structure and Function, 222(3), 1481–1493. doi: 10.1007/s00429-016-1288-8

Cacucci, F., Lever, C., Wills, T. J., Burgess, N., & O’Keefe, J. (2004). Theta-modulated place-by-direction cells in the hippocampal formation in the rat. Journal of Neuroscience, 24 (38), 8265–8277. doi: 10.1523/JNEUROSCI.2635-04.2004

Carp, J., Park, J., Hebrank, A., Park, D. C., & Polk, T. A. (2011). Age-Related Neural Dedifferentiation in the Motor System. PLoS ONE, 6(12), e29411. doi: 10.1371/journal.pone.0029411

Carp, J., Park, J., Polk, T. A., & Park, D. C. (2011). Age differences in neural distinctiveness revealed by multi-voxel pattern analysis. NeuroImage, 56(2), 736–743. doi: 10.1016/j.neuroimage.2010.04.267

Chersi, F., & Burgess, N. (2015). The Cognitive Architecture of Spatial Navigation: Hippocampal and Striatal Contributions (Vol. 88) (No. 1). Cell Press. doi: 10.1016/j.neuron.2015.09.021

Chowdhury, R., Guitart-Masip, M., Lambert, C., Dayan, P., Huys, Q., Düzel, E., & Dolan, R. J. (2013). Dopamine restores reward prediction errors in old age. Nature Neuroscience, 16(5), 648–653. doi: 10.1038/nn.3364

Cohen, J. D., & Servan-Schreiber, D. (1992). Context, Cortex, and Dopamine: A Connectionist Approach to Behavior and Biology in Schizophrenia. Psychological Review, 99(1), 45–77. doi: 10.1037/0033-295X.99.1.45

Cools, R., & D’Esposito, M. (2011). Inverted-U-shaped dopamine actions on human working memory and cognitive control (Vol. 69) (No. 12). Elsevier. doi: 10.1016/j.biopsych.2011.03.028

Cox, R. W., & Hyde, J. S. (1997). Software tools for analysis and visualization of fmri data. NMR in Biomedicine, 10(4-5), 171–178. doi: 10.1002/(SICI)1099-1492(199706/08)10:4/5¡171::AID-NBM453¿3.0.CO;2-L

Dale, A. M., Fischl, B., & Sereno, M. I. (1999). Cortical surface-based analysis: I. segmentation and surface reconstruction. NeuroImage, 9(2), 179–194. doi: 10.1006/nimg.1998.0395

El-Ghundi, M., Fletcher, P. J., Drago, J., Sibley, D. R., O’Dowd, B. F., & George, S. R. (1999). Spatial learning deficit in dopamine D1 receptor knockout mice. European Journal of Pharmacology, 383(2), 95–106. doi: 10.1016/S0014-2999(99)00573-7

Esteban, O., Birman, D., Schaer, M., Koyejo, O. O., Poldrack, R. A., & Gorgolewski, K. J. (2017). MRIQC: Advancing the automatic prediction of image quality in MRI from unseen sites. PLoS ONE, 12(9), e0184661. doi: 10.1371/journal.pone.0184661

Esteban, O., Blair, R., Markiewicz, C. J., Berleant, S. L., Moodie, C., Ma, F., … Gorgolewski, K. J. (2018). fmriprep. Software. doi: 10.5281/zenodo.852659

Esteban, O., Markiewicz, C., Blair, R. W., Moodie, C., Isik, A. I., Erramuzpe Aliaga, A., … Gorgolewski, K. J. (2018). fMRIPrep: a robust preprocessing pipeline for functional MRI. Nature Methods. doi: 10.1038/s41592-018-0235-4

Flossmann, T., & Rochefort, N. L. (2021). Spatial navigation signals in rodent visual cortex (Vol. 67). Elsevier Ltd. doi: 10.1016/j.conb.2020.11.004

Fonov, V., Evans, A., McKinstry, R., Almli, C., & Collins, D. (2009). Unbiased nonlinear average age-appropriate brain templates from birth to adulthood. NeuroImage, 47, Supplement 1, S102. doi: 10.1016/S1053-8119(09)70884-5

Garrett, D. D., Nagel, I. E., Preuschhof, C., Burzynska, A. Z., Marchner, J., Wiegert, S., … Lindenberger, U. (2015). Amphetamine modulates brain signal variability and working memory in younger and older adults. Proceedings of the National Academy of Sciences of the United States of America, 112 (24), 7593–7598. doi: 10.1073/pnas.1504090112

Goodridge, J. P. (1998). Cue control and head direction cells. Behavioral Neuroscience, 112(4), 749. doi: 10.1037/0735-7044.112.4.749

Gorgolewski, K., Burns, C. D., Madison, C., Clark, D., Halchenko, Y. O., Waskom, M. L., & Ghosh, S. (2011). Nipype: a flexible, lightweight and extensible neuroimaging data processing framework in python. Frontiers in Neuroinformatics, 5, 13. doi: 10.3389/fninf.2011.00013

Gorgolewski, K., Esteban, O., Markiewicz, C. J., Ziegler, E., Ellis, D. G., Notter, M. P., … Ghosh, S. (2018). Nipype. Software. doi: 10.5281/zenodo.596855

Granado, N., Ortiz, O., Suárez, L. M., Martín, E. D., Ceña, V., Solís, J. M., & Moratalla, R. (2008). D1 but not D5 dopamine receptors are critical for LTP, spatial learning, and LTP-induced arc and zif268 expression in the hippocampus. Cerebral Cortex, 18 (1), 1–12. doi: 10.1093/cercor/bhm026

Greve, D. N., & Fischl, B. (2009). Accurate and robust brain image alignment using boundary-based registration. NeuroImage, 48(1), 63–72. doi: 10.1016/j.neuroimage.2009.06.060

Guitchounts, G., Masís, J., Wolff, S. B., & Cox, D. (2020). Encoding of 3D Head Orienting Movements in the Primary Visual Cortex. Neuron, 108(3), 512–525.e4. doi: 10.1016/j.neuron.2020.07.014

Jeffery, K. J. (2007). Integration of the sensory inputs to place cells: What, where, why, and how? Hippocampus, 17(9), 775–785. doi: 10.1002/HIPO.20322

Jenkinson, M., Bannister, P., Brady, M., & Smith, S. (2002). Improved optimization for the robust and accurate linear registration and motion correction of brain images. NeuroImage, 17(2), 825–841. doi: 10.1006/nimg.2002.1132

Kentros, C. G., Agnihotri, N. T., Streater, S., Hawkins, R. D., & Kandel, E. R. (2004). Increased attention to spatial context increases both place field stability and spatial memory. Neuron, 42(2), 283–295. doi: 10.1016/S0896-6273(04)00192-8

Klein, A., Ghosh, S. S., Bao, F. S., Giard, J., Häme, Y., Stavsky, E., … Keshavan, A. (2017). Mindboggling morphometry of human brains. PLoS Computational Biology, 13(2), e1005350. doi: 10.1371/journal.pcbi.1005350

Kobelt, M., Sommer, V. R., Keresztes, A., Werkle-Bergner, M., & Sander, M. C. (2021). Tracking Age Differences in Neural Distinctiveness across Representational Levels. The Journal of Neuroscience, 41 (15), 3499–3511. doi: 10.1523/jneurosci.2038-20.2021

Koch, C., Li, S.-C., Polk, T. A., & Schuck, N. W. (2020). Effects of aging on encoding of walking direction in the human brain. Neuropsychologia, 141, 107379. doi: 10.1016/j.neuropsychologia.2020.107379

Koen, J. D., Hauck, N., & Rugg, M. D. (2019). The relationship between age, neural differentiation, and memory performance. Journal of Neuroscience, 39(1), 149–162. doi: 10.1523/JNEUROSCI.1498-18.2018

Kravitz, D. J., Saleem, K. S., Baker, C. I., & Mishkin, M. (2011). A new neural framework for visuospatial processing. Nature Reviews Neuroscience, 12 (4), 217–230. doi: 10.1038/nrn3008

Kroemer, N. B., Lee, Y., Pooseh, S., Eppinger, B., Goschke, T., & Smolka, M. N. (2019). L-DOPA reduces model-free control of behavior by attenuating the transfer of value to action. NeuroImage, 186, 113–125. doi: 10.1016/j.neuroimage.2018.10.075

Lanczos, C. (1964). Evaluation of noisy data. Journal of the Society for Industrial and Applied Mathematics Series B Numerical Analysis, 1 (1), 76–85. doi: 10.1137/0701007

Lenth, R. V. (2021). emmeans: Estimated marginal means, aka least-squares means [Computer software manual]. (R package version 1.6.1)

Lester, A. W., Moffat, S. D., Wiener, J. M., Barnes, C. A., & Wolbers, T. (2017). The Aging Navigational System. Neuron, 95(5), 1019–1035. doi: 10.1016/J.NEURON.2017.06.037

Leventhal, A. G., Wang, Y., Pu, M., Zhou, Y., & Ma, Y. (2003). GABA and its agonists improved visual cortical function in senescent monkeys. Science (New York, N.Y.), 300(5620), 812–5. doi: 10.1126/science.1082874

Li, S. C., Lindenberger, U., & Bäckman, L. (2010). Dopaminergic modulation of cognition across the life span. Neuroscience and Biobehavioral Reviews, 34 (5), 625–630. doi: 10.1016/j.neubiorev.2010.02.003

Li, S. C., Lindenberger, U., Hommel, B., Aschersleben, G., Prinz, W., & Baltes, P. B. (2004). Transformations in the Couplings Among Intellectual Abilities and Constituent Cognitive Processes Across the Life Span. Psychological Science, 15 (3), 155–163. doi: 10.1111/j.0956-7976.2004.01503003.x

Li, S.-C., Lindenberger, U., & Sikström, S. (2001). Aging cognition: from neuromodulation to representation. Trends in Cognitive Sciences, 5(11), 479–486. doi: 10.1016/S1364-6613(00)01769-1

Li, S. C., Papenberg, G., Nagel, I. E., Preuschhof, C., Schröder, J., Nietfeld, W., … Bäckman, L. (2013). Aging magnifies the effects of dopamine transporter and D2 receptor genes on backward serial memory. Neurobiology of Aging, 34 (1), 358.e1–358.e10. doi: 10.1016/j.neurobiolaging.2012.08.001

Li, S.-C., & Rieckmann, A. (2014). Neuromodulation and aging: implications of aging neuronal gain control on cognition. Current Opinion in Neurobiology, 29, 148–158. doi: 10.1016/j.conb.2014.07.009

Liang, Z., Yang, Y., Li, G., Zhang, J., Wang, Y., Zhou, Y., & Leventhal, A. G. (2010). Aging affects the direction selectivity of MT cells in rhesus monkeys. Neurobiology of Aging, 31 (5), 863–873. doi: 10.1016/J.NEUROBIOLAGING.2008.06.013

Lidow, M. S., Goldman-Rakic, P. S., Rakic, P., & Innis, R. B. (1989). Dopamine D2 receptors in the cerebral cortex: distribution and pharmacological characterization with [3H]raclopride. Proceedings of the National Academy of Sciences, 86(16), 6412–6416. doi: 10.1073/PNAS.86.16.6412

Mazziotta, J. C., Toga, A. W., Evans, A., Fox, P., & Lancaster, J. (1995). A Probabilistic Atlas of the Human Brain: Theory and Rationale for Its Development: The International Consortium for Brain Mapping (ICBM). NeuroImage, 2 (2, Part A), 89–101. doi: 10.1006/nimg.1995.1012

McNamara, C. G., & Dupret, D. (2017). Two sources of dopamine for the hippocampus. Trends in Neurosciences, 40(7), 383–384. doi: 10.1016/j.tins.2017.05.005

Moffat, S. D. (2009). Aging and Spatial Navigation: What Do We Know and Where Do We Go? Neuropsychology Review, 19(4), 478–489. doi: 10.1007/s11065-009-9120-3

Owesson-White, C. A., Cheer, J. F., Beyene, M., Carelli, R. M., & Wightman, R. M. (2008). Dynamic changes in accumbens dopamine correlate with learning during intracranial self-stimulation. Proceedings of the National Academy of Sciences of the United States of America, 105(33), 11957–11962. doi: 10.1073/pnas.0803896105

Papenberg, G., Bäckman, L., Nagel, I. E., Nietfeld, W., Schröder, J., Bertram, L., … Li, S. C. (2014). COMT polymorphism and memory dedifferentiation in old age. Psychology and Aging, 29(2), 374–383. doi: 10.1037/a0033225

Park, D. C., Polk, T. A., Park, R., Minear, M., Savage, A., & Smith, M. R. (2004). From The Cover: Aging reduces neural specialization in ventral visual cortex. Proceedings of the National Academy of Sciences, 101 (35), 13091–13095. doi: 10.1073/pnas.0405148101

Pedregosa, F., Michel, V., Grisel, O., Blondel, M., Prettenhofer, P., Weiss, R., … Duchesnay, É. (2011). Scikit-learn: Machine Learning in Python. Journal of Machine Learning Research, 12, 2825–2830. doi: 10.1007/s13398-014-0173-7.2

Power, J. D., Mitra, A., Laumann, T. O., Snyder, A. Z., Schlaggar, B. L., & Petersen, S. E. (2014). Methods to detect, characterize, and remove motion artifact in resting state fmri. NeuroImage, 84(Supplement C), 320–341. doi: 10.1016/j.neuroimage.2013.08.048

R Core Team. (2021). R: A language and environment for statistical computing [Computer software manual]. Vienna, Austria.

Satterthwaite, T. D., Elliott, M. A., Gerraty, R. T., Ruparel, K., Loughead, J., Calkins, M. E., … Wolf, D. H. (2013). An improved framework for confound regression and filtering for control of motion artifact in the preprocessing of resting-state functional connectivity data. NeuroImage, 64 (1), 240–256. doi: 10.1016/j.neuroimage.2012.08.052

Schmolesky, M. T., Wang, Y., Pu, M., & Leventhal, A. G. (2000). Degradation of stimulus selectivity of visual cortical cells in senescent rhesus monkeys. Nature Neuroscience, 3(4), 384–390. doi: 10.1038/73957

Schuck, N. W., Doeller, C. F., Polk, T. A., Lindenberger, U., & Li, S. C. (2015).Human aging alters the neural computation and representation of space. NeuroImage, 117, 141–150. doi: 10.1016/j.neuroimage.2015.05.031

Schuck, N. W., Doeller, C. F., Schjeide, B.-M. M., Schröder, J., Frensch, P. A., Bertram, L., & Li, S.-C. (2013). Aging and KIBRA/WWC1 genotype affect spatial memory processes in a virtual navigation task. Hippocampus, 23(10), 919–930. doi: 10.1002/hipo.22148

Schuck, N. W., Petok, J. R., Meeter, M., Schjeide, B. M. M., Schröder, J., Bertram, L., … Li, S. C. (2018). Aging and a genetic KI-BRA polymorphism interactively affect feedback- and observation-based probabilistic classification learning. Neurobiology of Aging, 61, 36–43. doi: 10.1016/j.neurobiolaging.2017.08.026

Servan-Schreiber, D., Printz, H., & Cohen, J. (1990). A network model of catecholamine effects: gain, signal-to-noise ratio, and behavior. Science, 249(4971), 892–895. doi: 10.1126/science.2392679

Shine, J. P., Valdés-Herrera, J. P., Hegarty, M., & Wolbers, T. (2016). The human retrosplenial cortex and thalamus code head direction in a global reference frame. Journal of Neuroscience, 36(24), 6371–6381. doi: 10.1523/JNEUROSCI.1268-15.2016

Sommer, V. R., Fandakova, Y., Grandy, T. H., Shing, Y. L., Werkle-Bergner, M., & Sander, M. C. (2019). Neural Pattern Similarity Differentially Relates to Memory Performance in Younger and Older Adults. The Journal of Neuroscience, 39(41), 8089–8099. doi: 10.1523/JNEUROSCI.0197-19.2019

Spiers, H. J., & Barry, C. (2015). Neural systems supporting navigation. Current Opinion in Behavioral Sciences, 1, 47–55. doi: 10.1016/j.cobeha.2014.08.005

St-Laurent, M., Abdi, H., Bondad, A., & Buchsbaum, B. R. (2014). Memory reactivation in healthy aging: Evidence of stimulus-specific dedifferentiation. Journal of Neuroscience, 34 (12), 4175–4186. doi: 10.1523/JNEUROSCI.3054-13.2014

Taube, J. S. (2007). The Head Direction Signal: Origins and Sensory-Motor Integration. Annual Review of Neuroscience, 30(1), 181–207. doi: 10.1146/annurev.neuro.29.051605.112854

Taube, J. S., Muller, R. U., & Ranck, J. B. (1990). Head-direction cells recorded from the postsubiculum in freely moving rats. II. Effects of environmental manipulations. Journal of Neuroscience, 10(2), 436–447. doi: 10.1523/jneurosci.10-02-00436.1990

Thurley, K., Senn, W., & Lüscher, H. R. (2008). Dopamine increases the gain of the input-output response of rat prefrontal pyramidal neurons. Journal of Neurophysiology, 99(6), 2985–2997. doi: 10.1152/jn.01098.2007

Thurm, F., Schuck, N. W., Fauser, M., Doeller, C. F., Stankevich, Y., Evens, R., … Li, S.-C. (2016). Dopamine modulation of spatial navigation memory in Parkinson’s disease. Neurobiology of Aging, 38, 93–103. doi: 10.1016/j.neurobiolaging.2015.10.019

Tustison, N. J., Avants, B. B., Cook, P. A., Zheng, Y., Egan, A., Yushkevich, P. A., & Gee, J. C. (2010). N4itk: Improved n3 bias correction. IEEE Transactions on Medical Imaging, 29(6), 1310–1320. doi: 10.1109/TMI.2010.2046908

Vijayraghavan, S., Wang, M., Birnbaum, S. G., Williams, G. V., & Arnsten, A. F. (2007). Inverted-U dopamine D1 receptor actions on prefrontal neurons engaged in working memory. Nature Neuroscience, 10(3), 376–384. doi: 10.1038/nn1846

Volkow, N. D., Gur, R. C., Wang, G. J., Fowler, J. S., Moberg, P. J., Ding, Y. S., … Logan, J. (1998). Association between decline in brain dopamine activity with age and cognitive and motor impairment in healthy individuals. American Journal of Psychiatry, 155(3), 344–349. doi: 10.1176/ajp.155.3.344

Wickham, H. (2016). ggplot2: Elegant graphics for data analysis. Springer-Verlag New York.

Wolbers, T., Dudchenko, P. A., & Wood, E. R. (2014). Spatial memory - A unique window into healthy and pathological aging (Vol. 6) (No. MAR). Frontiers Media SA. doi: 10.3389/fnagi.2014.00035

Zhang, Y., Brady, M., & Smith, S. (2001). Segmentation of brain MR images through a hidden markov random field model and the expectation-maximization algorithm. IEEE Transactions on Medical Imaging, 20(1), 45–57. doi: 10.1109/42.906424

